# Exosomal melanotransferrin as a potential biomarker for metastatic melanoma

**DOI:** 10.1101/2020.11.09.373852

**Authors:** Anne Bonhoure, Laurent Henry, Marie Morille, Nesrine Aissaoui, Gaëtan Bellot, Pierre-Emmanuel Stoebner, Michel Vidal

## Abstract

Cutaneous melanoma is the most lethal type of skin cancer. Early detection is crucial to improve the outcome of melanoma patients. The identification of noninvasive prognostic biomarkers for the follow-up of melanoma patients is still in demand for clinical use. We show here that exosomal melanotransferrin (MTf) fulfils the biomarker characteristics required to meet this demand. MTf is typically overexpressed in melanoma cells compared to other cell types – including cancer cells – and is efficiently sorted and secreted with nanovesicles, or so-called exosomes, due to its membrane-anchoring by a glycosylphosphatidylinositol. MTf is exposed on the surface of exosomes and is accessible for antibody recognition. An ELISA was set-up to quantify MTf after immobilization of nanovesicles through the exosomal constituent tetraspanins CD63. MTf was detected using low number of exosomes purified from melanoma cell line cultures, and MTf detection was abolished by phosphatidylinositol-specific phospholipase C (PI-PLC) treatment. This exosomal MTf ELISA was able to discriminate an equal number of assayed exosomes purified from two different melanoma cell lines (A-375 vs SK-MEL-28). Moreover, plasma samples from patients with melanoma and noncancer disease were assayed using this ELISA and exosomal MTf was validated as a potential melanoma biomarker.

## Introduction

Melanoma is a very aggressive cancer whose incidence is rapidly increasing worldwide. According to the American Institute for Cancer Research, nearly 300,000 new cases were diagnosed worldwide in 2018. Invasive melanoma incidence is still rising faster than that of any other cancer despite the advent of novel molecularly targeted and immune therapies^1,2^. Clinical evidence available for prognostic assessment of patients with metastatic melanoma remains insufficient despite extensive research on the identification of effective markers. The only serum biomarker validated by the American Joint Committee on Cancer (AJCC) to provide melanoma prognostic information is the lactate dehydrogenase (LDH) level in blood. Serum levels of other markers such as melanoma inhibitory activity protein (MIA)^3^, S100 calcium binding protein (S100B)^4^ or plasmatic 20S proteasome (p-proteasome)^5^ are, like LDH, correlated with disease progression, but they are not more specific to melanoma cell proliferation. There is thus a crucial need for developing new specific candidate biomarkers to enhance melanoma diagnosis and therapeutic monitoring^6,7^. Demand is therefore currently high for peripheral blood biomarkers because of their ready accessibility for analysis.

Extracellular vesicles (EVs) are such peripheral blood components and are nowadays being actively studied for their biomarker potential. Exosomes, a subset of EVs, are small extracellular vesicles (approx. 80 nm) originating from the endosomal compartment of cells that are generated by membrane budding toward the endosomal lumen^8^. These intraluminal vesicles (ILVs) are released in the extracellular space upon fusion of multivesicular endosomes (MVEs) with the plasma membrane^9,10^. The protein composition of exosomes found in body fluids is related to the cells that produce them. Several proteins, e.g. Tsg101, CD63, participate in the biogenesis of ILVs and are constitutively found in exosomal vesicles, while some cargo proteins are specific to the producing cell. Moreover, exosomes contain nucleic acids, including miRNAs, which further enhances their interest as potential biomarkers^11,12^.

The main aim of our study was to characterize a protein that is: (1) specific to melanoma cells, (2) efficiently sorted in exosomes secreted by these cells, and (3) exposed on the surface of vesicles. Our previous studies on exosomes secreted by reticulocytes revealed active sorting of glycosylphosphatidylinositol (GPI)-anchored proteins in exosomes^13,14^. Otherwise, melanotransferrin (MTf) is one of the first cellular markers of melanoma that was identified in the early 1980s^15^. Interestingly, MTf is a GPI-anchored protein^16^ shown to be overexpressed in melanoma cells as compared to melanocytes^15,17^. We thus studied MTf as a putative marker protein that could be exposed on the surface of melanoma exosomes. We found that MTf is enriched in secreted exosomes as compared to cell lysates of melanoma cell lines (A-375, SK-MEL-28). In contrast, MTf has not been found in exosomes released by other types of cancer cell lines, including MCF7, Caco2, K562 and HeLa cells. GPI-anchoring of MTf on the exosomal membrane was demonstrated by Triton X-114 phase partitioning and susceptibility to phosphatidylinositol-specific phospholipase C (PI-PLC). Moreover, by vesicle immunoisolation we demonstrated that CD63 and MTf were both present on the surface of exosomes secreted by melanoma cell lines. We thus set up an immunocapture-based ELISA to detect MTf on immobilized CD63-positive exosomes, first collected from the supernatant of melanoma cell cultures and then directly using plasma from patients. Here we provide evidence that exosomal melanotransferrin could be used as melanoma biomarker.

## Materials and methods

### Materials

Triton X-114 (Sigma), bromophenol blue (Bio-Rad), *Bacillus cereus* PI-PLC (Molecular Probe, P6466), Clarity Max™ Western ECL substrates (Bio-Rad) were used. Prestained molecular markers were from Thermo Scientific (PageRuler) or VWR (PeqGOLD marker V). Immobilon-P transfer membranes were from Millipore. Dynabeads™ Protein G were obtained from Life Technologies (10003D). Gelatin Type B from bovine skin was from Sigma (G-9382). Protease inhibitor cocktail (11697498001) was from Roche. 3,3’,5,5’-Tetramethylbenzidine (TMB) ELISA peroxidase substrate (UP664781) was obtained from Interchim. Nunc MaxiSorp™ flat-bottom plates were from Thermo Fischer scientific (44-2404-21). Accutase™ solution was obtained from Gibco (A11105).

### Antibodies

The following primary antibodies were used in Western blotting experiments: anti-MTf (R&D systems, MAB8175), CD63 (Abcam, ab134045), Tsg101 (Abcam, ab83), Alix (Abcam, ab186429), Flotillin-1 (BD Biosciences, clone 18), hsc70 (Stressgen, 1B5), CD9 (GeneTex, GTX62294), Lyn (Santa Cruz, sc-7274), syntenin-1 (GeneTex, GTX108470), CEACAM6 (Temecula, MABT 323). HRP-conjugated secondary antibodies used for Western blot and ELISA were purchased from Jackson ImmunoResearch Laboratories.

Mouse monoclonal anti-MTf (R&D Systems, MAB81751, dilution 1/50) antibodies were used for flow cytometry and immunofluorescence using anti-mouse IgG conjugated with Alexa Fluor 488 (Invitrogen, A11029, dilution 1/2000) as secondary antibodies. Cholera toxin B-chain conjugated with Alexa Fluor 555 was from Thermo Fischer scientific (C-34775).

Anti-CD63 (Abcam, ab8219) was used for exosome immunoisolation. Anti-CD63 (Abcam, ab252919) and anti-MTf (R&D systems, MAB81751 or Abcam, ab201922) antibodies were used for immunocapture-based ELISA.

### Cells

Melanoma A-375 and SK-MEL-28, breast adenocarcinoma MCF7, colon cancer Caco2, chronic myelogenous leukemia K562 and cervix adenocarcinoma HeLa cell lines were maintained in DMEM containing 10% heat inactivated FBS, 2 mM GlutaMax-I™, 1 mM sodium pyruvate, 0.1 mM non-essential amino acids, 100 UI/ml penicillin and 100 µg/ml streptomycin.

Exosome production was carried out for 24 h in DMEM containing 1% EV-depleted FBS^18^ to limit serum protein retrieval during exosome purification, after two DMEM washes of the cells. For western blot experiments, adherent cells were washed once in PBS before trypsinization. Cell pellets were washed twice in ice-cold PBS and lysed in RIPA (150 mM NaCl, 20 mM Tris pH 7.4, 1 mM EDTA, 1% DOC, 1% NP40, 2 mM DTT and antiprotease cocktail) for 20 min on ice. After centrifugation (20 min at 20 000 x g), the protein concentration in supernatants was determined using Coomassie protein assay reagent (1856209, Thermo Scientific) and bovine serum albumin as standard.

### Plasma of patients

Plasma of melanoma patients was obtained from MelBase, a national cohort of melanoma stage IV and unresectable stage III patients (https://ichgcp.net/clinical-trials-registry/NCT02828202). Blood samples from patients with psoriasis were obtained after signature of a consent form, following approval of the procedure by the local ethics committee in accordance with French regulations. Plasma was assayed for plasmatic p-proteasome, as previously described^5^.

### Indirect flow cytometry

A-375 and SK-MEL-28 cells were washed in PBS, detached by accutase™ treatment and washed again in PBS (300 g, 5 min, 4°C). The whole labelling procedure was carried out at 4°C : 1.2 10^6^ cells were incubated for 1h in 100 µl 3% BSA/PBS containing (or not) anti-MTf antibody. After three washes with 3.5 ml BSA/PBS, cells were incubated in the dark for 30 min in 100 µl BSA/PBS containing (or not) secondary antibody. Cells were washed twice in BSA/PBS, then once in PBS, and pellets were resuspended in 600 µl PBS. Fluorescence was measured in the FL1 channel of a BD FACSCanto cytometer (BD FACSDiva software v8.0.2) with FITC gain kept constant at 401 V all along acquisition. All data were subsequently analyzed using FlowJo v10.5.3 (FlowJo, LLC, Ashland OR).

### Immunofluorescence imaging

Cells grown on glass coverslips were washed three times in cold DMEM and incubated for 30 min at 4°C with cholera toxin B-chain conjugated with Alexa Fluor 555 (1/1000 in DMEM). After washing with DMEM, cells were fixed in 4% formaldehyde for 20 min at 4°C. After neutralization (10 min RT in PBS containing 50 mM NH_4_Cl) and washes, cells were blocked for 20 min in 0.1% BSA in PBS before 1 h incubation at RT with anti-MTf antibodies, followed by 3 washes in PBS/BSA and incubation with anti-mouse IgG Alexa Fluor 488 for 45 min. Nuclei were counter-stained with Hoechst and coverslips were mounted using Thermo Scientific™ Shandon™ immu-mount™.

Cells were observed using a ZEISS LSM880 confocal microscope equipped with diode (405 nm) and Argon (488 and 514 nm) excitation sources, a 63x (1.4 Oil Plan-apochromat) objective, and a GaAsP detector. ZEN software was used for image acquisitions.

To evidence difference in MTf expression between the two melanoma cell lines, images were first acquired with identical settings. Then, the laser power was lowered when observing SK-MEL-28 to visualize MTf labeling without pixel saturation. Controls (without 1^st^ antibody) were observed with the respective laser settings.

### Exosome purification

Exosomes were isolated by differential centrifugation as described previously^18^. Briefly, culture supernatants were successively centrifuged (10 min at 500 x g; 20 min at 20 000 x g; 3 h at 100 000 x g) and the final pellets were resuspended and washed once in PBS for use as crude pellet. For further purification, these crude pellets were resuspended, layered on top of a sucrose gradient (4 ml of 0.25 to 2 M sucrose, 20 mM HEPES/NaOH, pH 7.4) in a Beckman SW55 rotor (Beckman Coulter, Fullerton, CA) and centrifuged for 20 h at 45 000 rpm, after which 350 µl fractions were collected from the top of the tube using a peristaltic pump. For each fraction, 15 µl were used for determination of the refractive index and density, and 12 µl for Western blot analysis. Fractions positive for exosome markers and negative for 20S proteasomes, with densities ranging from 1.10 to 1.15 were pooled and washed in PBS (Beckman Ti50, 3 h at 42 000 rpm).

### Cryo-electron microscopy

For cryo-EM experiments, 3 μl of samples were applied to glow-discharged Quantifoil R 0.6/1 and R 1.2/1.3 grids (Quantifoil Micro tools GmbH, Germany), blotted for 3 s and then flash frozen in liquid ethane using the automatic plunge freezer EM GP2 at 100% relative humidity. All observations of EM-grids were carried out on a JEOL 2200FS FEG operating at 200 kV equipped with a 4k x 4k slow-scan CDD camera (Gatan inc.) under low-dose conditions (total dose of 20 electrons/Å^2^) in the zero-energy-loss mode with a slit width of 20 eV. Images were taken at a nominal magnification of 50 000 X, with defocus ranging from 0.6 to 2.5 μm.

### Particle size distribution

Samples from the purified exosomal fractions were examined by nanoparticle tracking analysis (NTA), as previously described^18^. Briefly, 1 ml of exosomes diluted in PBS (1/1000-1/2000, corresponding to a 1.10^7^ to 1.10^9^ NP/ml concentration range, as recommended by the manufacturer) was loaded using a syringe-pump holder (speed: 40 AU) into the sample chamber of a Nanosight NS300 (Malvern) equipped with a 405 nm laser. Acquisitions were obtained at 25°C with a sCMOS camera (level 15) to capture videos. Data analysis was performed with NTA 3.2 software (Nanosight) with a detection threshold of 5. The graphs represent the average finite track-length adjusted (FTLA) plot of the concentration (particle/ml) as a function of the particle size (nm) averaged from three captures (1 min each). The mean and mode of particle size distribution are indicated.

### Western blotting

Samples were separated by SDS-PAGE (10% or 12% acrylamide), and analyzed by Western blot, as previously described using enhanced chemiluminescence substrates and a Bio-Rad Chemidoc™ device^18^. PVDF membranes were stripped by incubation in 0.2 M glycine, 0.1% SDS, 1% Tween 20, pH 2.2, for 1 h at 50°C, and then washed, blocked and re-probed^19^.

When indicated, the protein expression levels were quantified from unsaturated images using Fiji software (NIH).

### Analysis of exosomal melanotransferrin anchoring

#### TX-114 partitioning

A stock solution of 12% Triton X-114 containing a trace of bromophenol blue was obtained by precondensation as described^20^. Exosome suspension (25 µg in 100 µl PBS) was mixed with 20 µl Triton X-114 stock solution to achieve a 2% final detergent concentration and incubated for 15 min at 4° C with double vortexing. Phase separation was obtained by incubation at 37° C for 30 min. The upper aqueous phase (Aφ) was withdrawn while the detergent phase was treated similarly twice after the addition of 100 µl Tris buffer saline (150 mM NaCl, 20 mM Tris pH 7.4) to obtain the final detergent phase (Dφ). Proteins contained in the two phases (Aφ and Dφ) were precipitated by cold acetone, resuspended in Laemmli buffer, loaded on 12% SDS-PAGE and analyzed by Western blot for the indicated proteins.

#### PI-PLC sensitivity

Exosomes purified from SK-MEL-28 (5 µg prot) were incubated or not with phosphatidylinositol-specific phospholipase C from *Bacillus cereus* (1 u) for 30 min at room temperature. Exosomes were then pelleted (TLA-100 rotor Beckman, 1 h at 90 000 rpm). Supernatants and pellets were then loaded on 12% SDS-PAGE and the indicated proteins were analyzed by Western blot.

### Immunoisolation of exosomes through CD63

Exosomes (25 μg prot) from A-375 cells were incubated or not with 5 µl anti-CD63 antibodies (Abcam, ab8219) for 2 h at room temperature. Dynabeads™ protein G (10 μl of 50% PBS slurry) was then added for 3 h at room temperature. After magnet separation, half of the supernatants (Sup), as well as the beads washed twice by PBS (beads), were mixed with Laemmli buffer, boiled and run on 10% SDS-PAGE before transfer and analysis by Western blot for the indicated proteins.

### Gelatin zymography

Exosomes and cell lysates (10 µg prot) from A-375 and SK-MEL-28 melanoma cell lines were mixed with non-reducing sample buffer, loaded and run at 150 V on a 7.5% SDS-PAGE containing 1 mg/ml gelatin. Gels were soaked (2 x 30 min) in washing buffer (2.5% Triton X-100, 50 mM Tris pH 7.5, 5 mM CaCl_2_, 1 µM ZnCl_2_) to remove SDS and incubated for 24 h at 37°C in incubation buffer (same as washing buffer but with 1% Triton X-100 instead of 2.5%). The gels were then stained with Coomassie blue and destained until bands corresponding to areas of enzyme activity could clearly be seen.

### Immunocapture-based ELISA (Sup figure 1A)

Wells of a 96-well plate (Nunc Maxisorb) were coated with 100 ng of anti-CD63 antibody (ab252919, Abcam) in 100 µl carbonate buffer (pH 9.6) overnight at 4°C with gentle agitation, washed twice with PBS and blocked by PBS-5% BSA for 2 h at room temperature (RT). Plates were then washed three times with PBS, and 100 µl volume of exosomes – or plasma diluted 1/20 in PBS – were incubated for 2 h at RT. After five washes with PBS, the wells were added with anti-MTf antibody (50 ng of MAB81751, R&D Systems) in PBS-5% BSA for 2 h at RT, washed again five times with PBS, and anti-mouse IgG-HRP (Jackson Immunoresearch) – diluted (1/5000) in PBS-5% BSA – was incubated for 1h at RT. After PBS washes, 100 µl TMB solution (UP664781) was added to the wells, color development was stopped by adding 50 µl H_2_SO_4_ (1 M) after 10-20 min incubation and absorbance was recorded at 450 nm using a TECAN Sunrise spectrophotometer. When indicated, exosomes were pre-incubated for 30 min at RT with 1% Triton X-100 or PI-PLC from *Bacillus cereus* (0.5 u).

**Figure 1:**
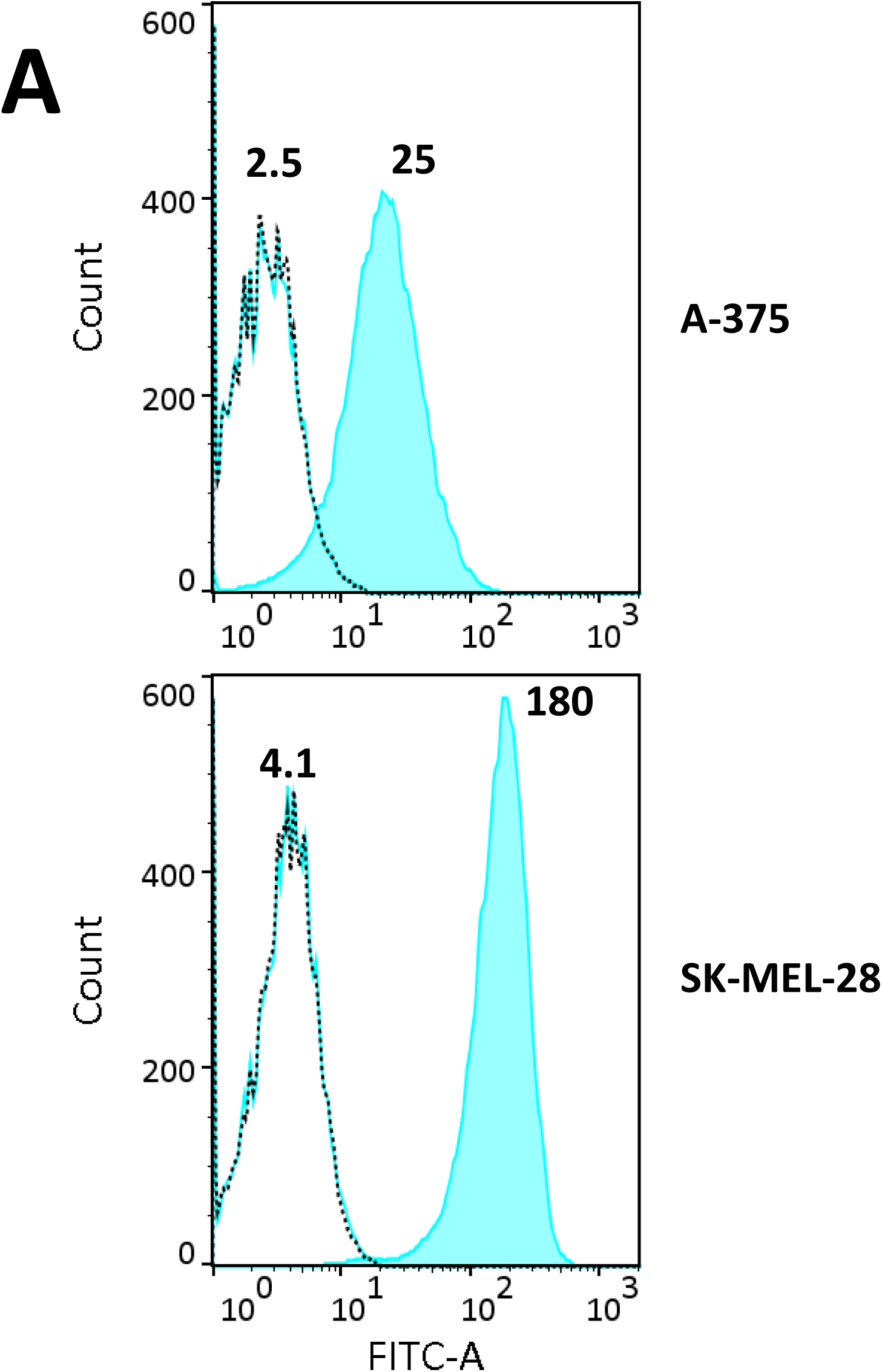
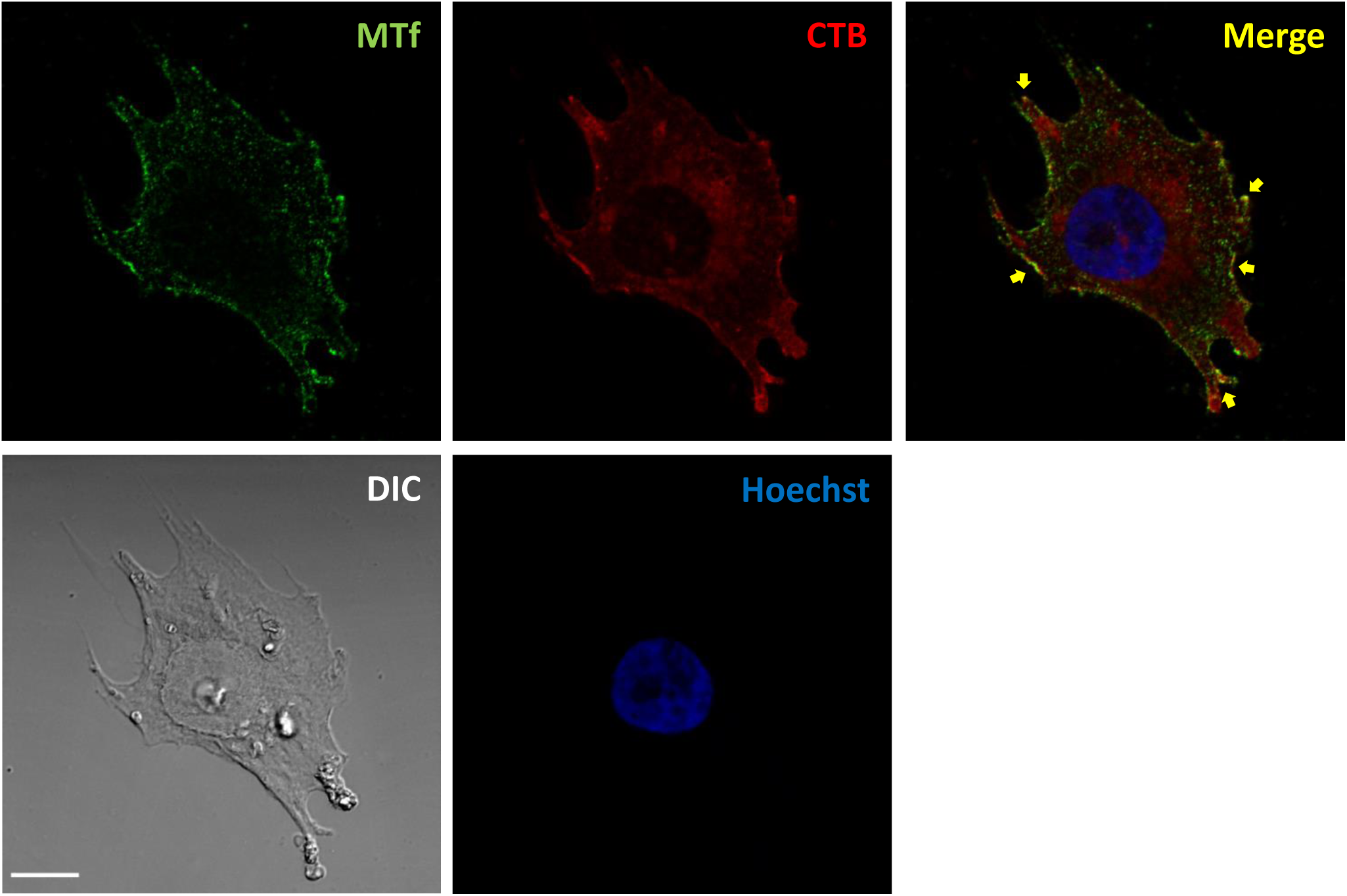
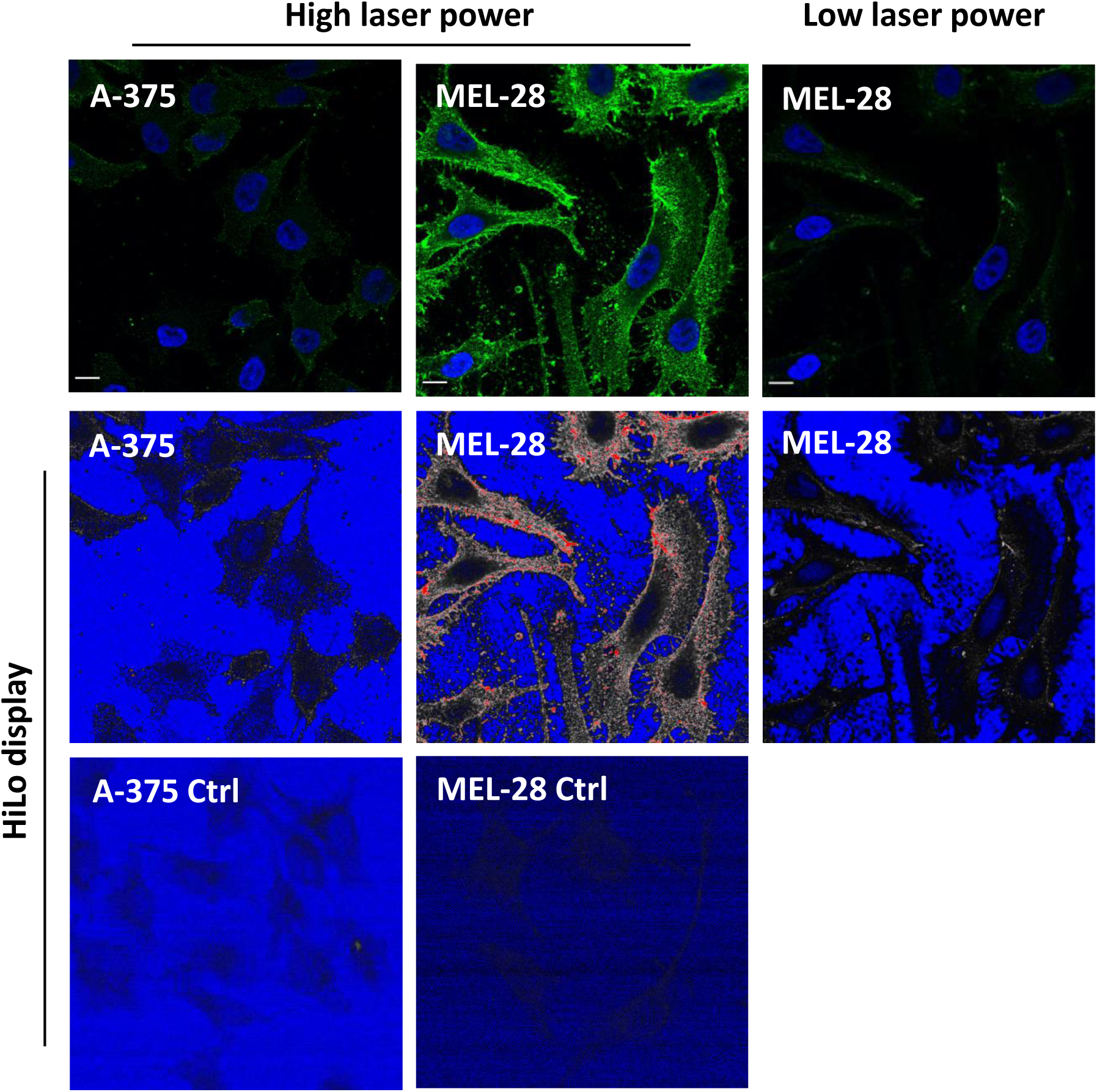
Analysis of the expression of melanotransferrin on the surface of melanoma cells. **A)** Flow cytometry analysis of MTf on the surface of A-375 and SK-MEL-28 cells was performed as described in the Materials and Methods. Histogram of fluorescence intensity showing cell autofluorescence (black dotted line), cells incubated with 2^nd^ antibody only (blue line) and cells fully immunolabeled (blue-filled pattern). Mean fluorescence intensities are indicated. **B)** A-375 cells were processed for immunofluorescence using anti-MTf antibody followed by anti-mouse IgG conjugated with Alexa Fluor 488 (green), and cholera toxin B chain conjugated with Alexa Fluor 555 (red). Images were acquired using a Confocal ZEISS LSM880 microscope (63X/1.4 oil). Brightness and contrast were adjusted using Fiji. Scale bar: 10 µm. **C)** A-375 and SK-MEL-28 cells were processed for MTf immunofluorescence, as above. Raw images are presented (higher panels). Laser intensity (1.0%) and gain (610) were first adjusted to image MTf on A-375 cells without pixel saturation, and SK-MEL-28 cells were acquired using the same settings (high laser power). Differential MTf expression is striking when pixel intensities are displayed in Fiji HiLo LUT. To avoid pixel saturation (red) on that same field, laser intensity (0.8%) and gain (500) had to be substantially lowered (low laser power). Scale bar: 10 µm.

## Results

### Melanotransferrin on the surface of melanoma cell lines

Two melanoma cell lines (A-375, SK-MEL-28) were used and respectively represented low and highly metastatic melanoma^21^, even though discrepancies were found with regard to this feature^22,23^. We first assessed melanotransferrin expression on the plasma membrane of the two cell lines by flow cytometry (Figure 1A). Note the difference of mean fluorescence intensity for A-375 cells (x10) and SK-MEL-28 (x44) compared to their respective negative controls, reflecting higher MTf expression on the SK-MEL-28 cell surface. Flow cytometry analysis on fixed cells – either permeabilized or not – gave similar results (not shown).

We also examined the pattern of MTf and ganglioside GM1 on the cell surface by confocal fluorescence microscopy. Since MTf is a GPI-anchored protein^24^, the possibility of colocalisation with GM1 in lipid rafts on the cell surface was investigated. Ganglioside GM1 was labeled through binding of cholera toxin B chain (CTB)-labeled with Alexa Fluor 555. Colocalisation of MTf and GM1 on the cell surface could be sparsely found in A-375 cells, as shown in Figure 1B. Strikingly, SK-MEL-28 cells were not labeled by fluorescent CTB compared to A-375 cells (not shown). The absence of ganglioside GM1 in SK-MEL-28 has already been described^25^, possibly because of ganglioside metabolism diverted to GD3 synthesis, influencing the malignant properties of melanoma cells^26^. In both cell lines, MTf presented a punctate (green) pattern, though stronger in SK-MEL-28 compared to A-375 cells when observed using the same laser setting (Figure 1C, left and middle panels). As shown, laser power was lowered for SK-MEL-28 observation to obtain similar pixel intensity compared to A-375 cells.

### Melanotransferrin is enriched in exosomes vs cell lysates of A375 and SK-MEL-28 cells

We assume that lipid rafts are membrane nanodomains that likely play a role during exosome biogenesis^27,28^. We have previously shown that lipid raft components such as GPI-anchored proteins are present in high amounts in exosomes secreted by reticulocytes^14,27^. Since melanotransferrin is a transferrin homolog that is found predominantly bound to the cell membrane via GPI anchoring^16,29^, we assessed the presence of MTf in exosomes secreted by A-375 and SK-MEL-28 melanoma cells. Exosomes from these two cell lines were purified and characterized, as previously described^18^. Cryo-EM experiments were performed before and after sucrose-gradient purification (Figure 2A). On crude ultracentrifugation pellet, we visualized a large variety of extracellular vesicles of various size and morphology (Figure 2A, left panel). Some of the vesicles are assigned into multilayer category with multiple vesicles inside a larger one with either spherical or elongated shape. In addition, vesicles debris and expelled contents are observed in the medium as indicator that sample contamination is dominating the sample. Sucrose gradient-purified exosomes show however more homogenous vesicles populations in size and morphology. Furthermore, contamination from medium proteins is not observed anymore (Figure 2A, right panel), which indicates the efficiency of the sucrose-gradient step for the isolation and purification of exosomes.

**Figure 2:**
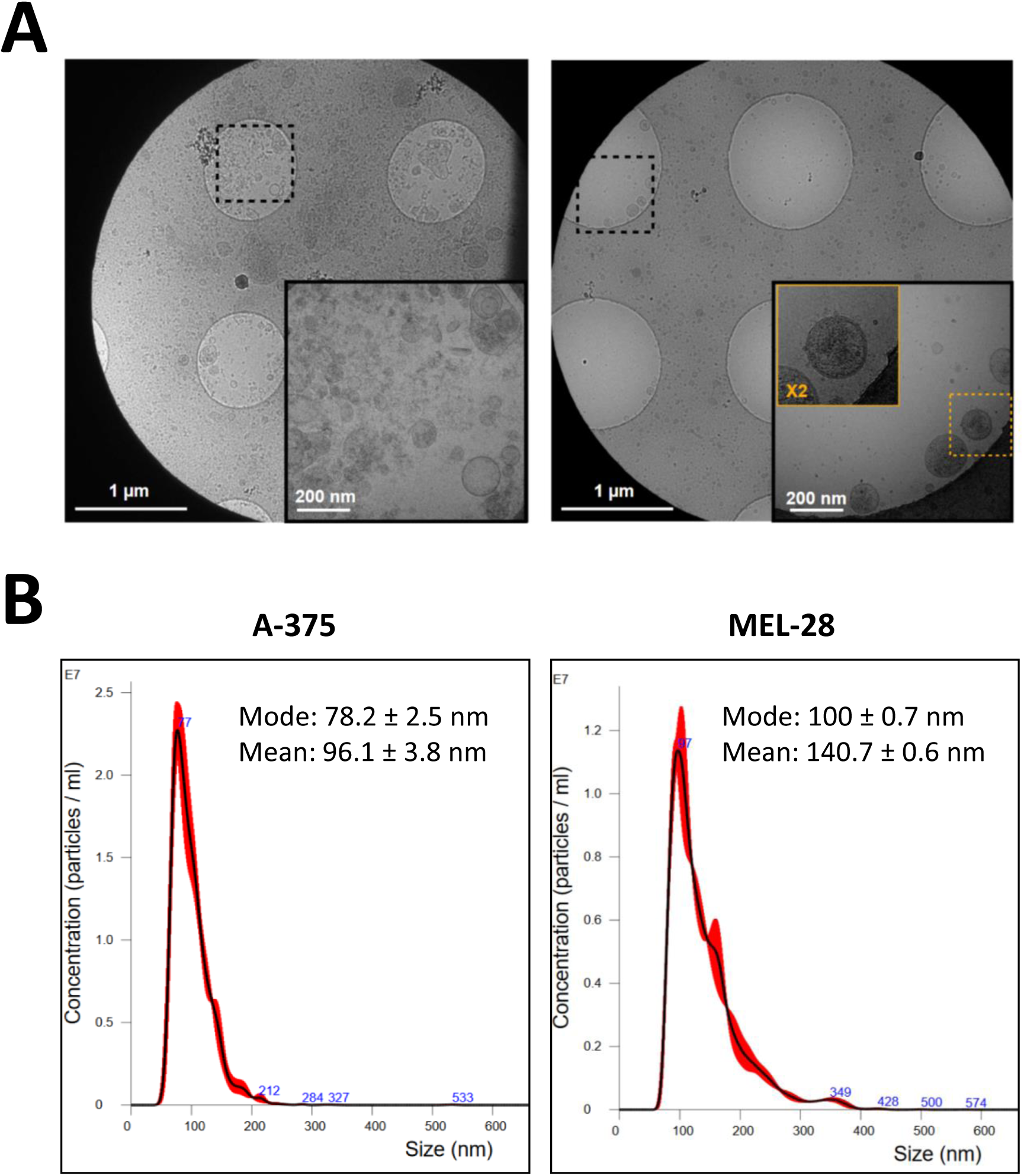
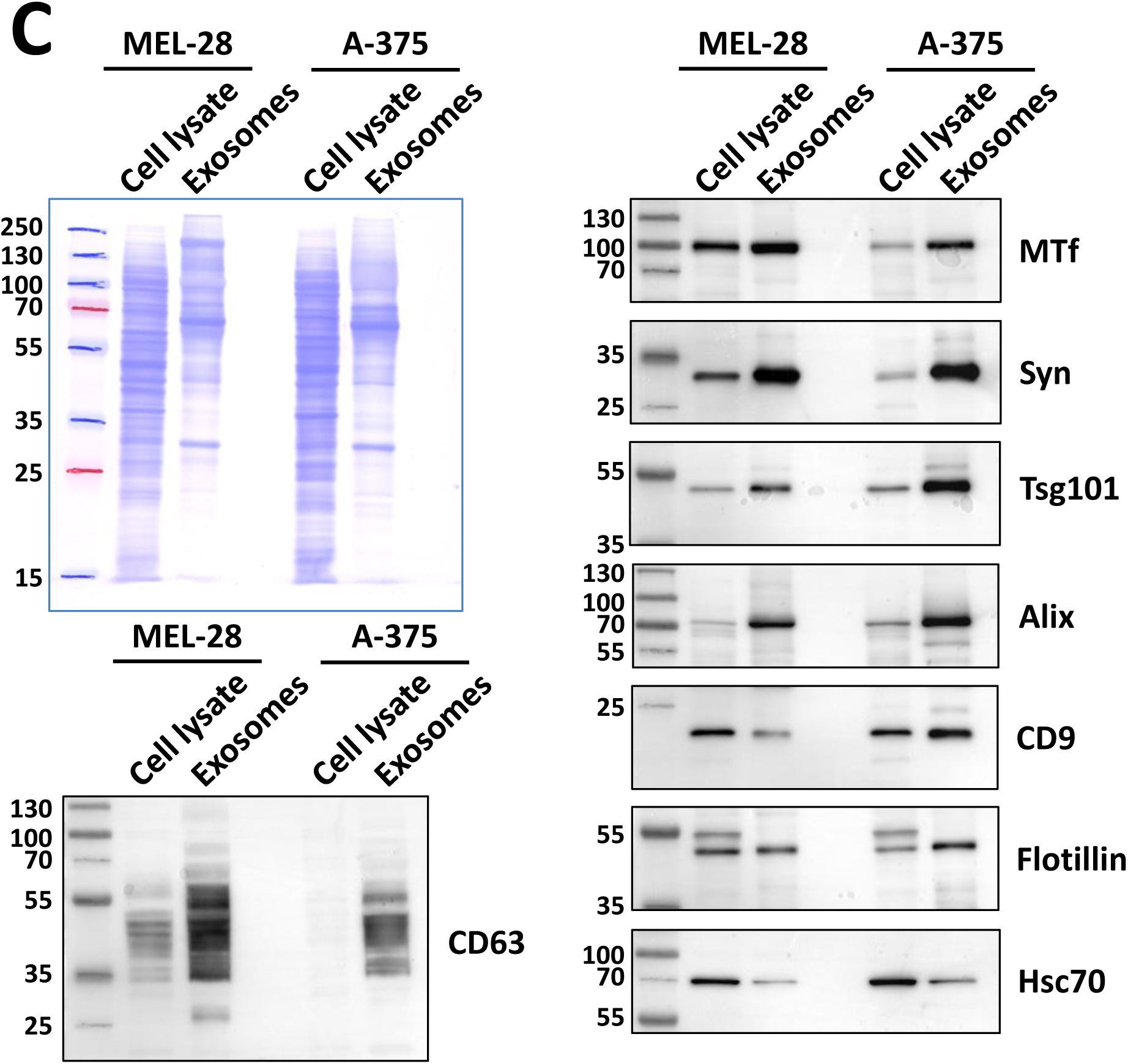
Purification and characterization of melanoma exosomes. SK-MEL-28 and A-375 melanoma cells were cultured for 24 h in medium containing 1% EV-depleted serum. Exosomes were purified from the culture medium as described in the Materials and Methods. **A)** Typical cryo-EM micrographs of crude 100,000 x g pellet (left panel) and exosomes further purified through sucrose gradient (right panel) imaged at different magnifications. **B)** Size distribution of purified exosome preparations from A-375 and SK-MEL-28 cells were analyzed by NTA. Mean and mode of particle size distributions are indicated. **C)** Cell lysates and purified exosomes (10 µg protein) were analyzed by Western blot for the indicated proteins after protein separation by 12% SDS-PAGE.

Accordingly, as assessed by nanoparticle tracking analysis (Figure 2B), the purified vesicles were highly monodispersed in size (80-100 nm), corresponding to the size expected for exosomes. As shown by Western blot, the purified vesicles contained *bona fide* markers – e.g. Alix, Tsg101, syntenin-1, CD63 – of exosomes^8^ (Figure 2C). Note especially their enrichment in exosomes compared to the same amount of protein from the cell lysate of the respective producing cells. When MTf was assessed by Western blot, enrichment in exosomes compared to cell lysates was observed for both cell lines regardless of the cellular MTf expression level (Figure 2C). This clearly indicated active sorting of MTf in exosomes. Note that MTf was more expressed in SK-MEL-28 compared to A-375 cells, in agreement with our flow cytometry and immunofluorescence analyses and with the findings of previous studies presenting SK-MEL-28 cells as the highest MTf-expressing cell line among a panel of other melanoma cell lines^15,30^. Note also the huge enrichment of CD63 in exosomes compared to the CD9 distribution in the two fractions – exosomes vs cell lysates – thus underlining the accuracy of CD63 compared to CD9 as exosomal markers and in agreement with their cellular localization according to the Human Protein Atlas database (www.proteinatlas.org).

### Melanotransferrin is associated with exosomes via a GPI-anchor

Since a soluble form of MTf has also been reported^31^, we examined the association of melanotransferrin with exosomes secreted by A-375 and SK-MEL-28 cells. A standard approach to identify GPI-anchored proteins is based on Triton X-114 phase partitioning assessment of their amphiphilic characteristics^20^. Following TX-114 treatment of exosomes secreted by both cell lines, we obtained a clear separation of transmembrane and amphiphilic proteins in the detergent phase, as compared to hydrophilic proteins in the aqueous phase (Figure 3). MTf was essentially found in the detergent phase – as well as tetraspanins CD9 and CD63 – thus supporting its glycolipid-anchoring on the exosomal membrane, whereas Tsg101, Alix and syntenin-1 were retrieved in the aqueous phase, as expected for cytosolic proteins. Note that the tyrosine kinase Lyn partitioned in both detergent and aqueous phases, which might reflect potential cleavage of its N-terminal domain which is critical for membrane association through double acylation, as we already described in exosomes from various cell lines^27^. Flotillin-1 was also partitioned in both detergent and aqueous phases, as already described^32^.

**Figure 3:**
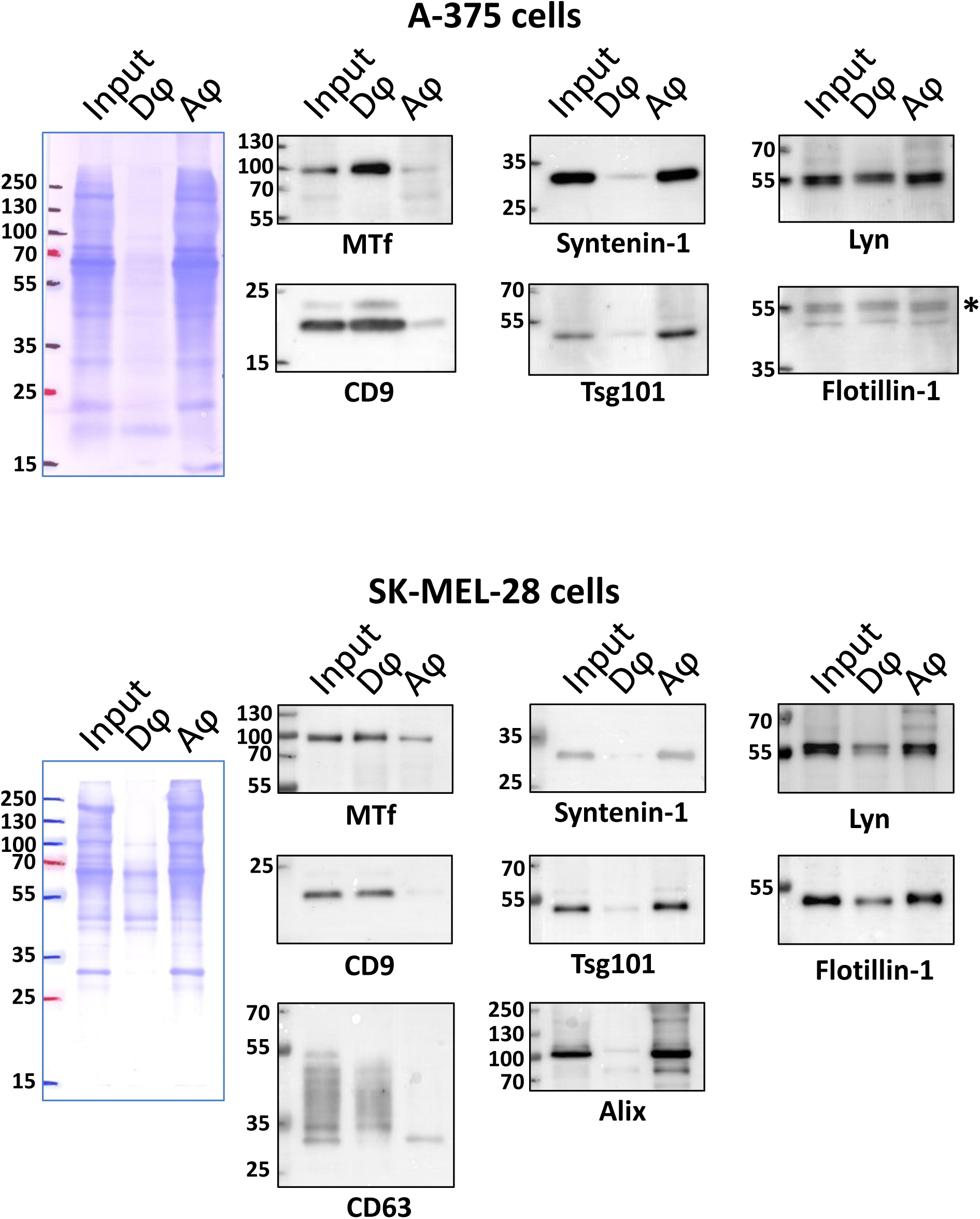
Analysis of exosomal melanotransferrin anchoring by TX-114 partitioning. Exosomes (25 µg) from A-375 (upper panel) or SK-MEL-28 cells (lower panel) were treated for TX-114 partitioning as described in the Materials and Methods. Detergent (Dφ) and aqueous (Aφ) phases were loaded on 12% SDS-PAGE after acetone precipitation and analyzed by Western blot for the indicated proteins. Input: 10 µg of respective exosomes. The asterisk (*) in the upper panel indicates bands remaining from previous Lyn blotting.

Another way to demonstrate GPI-anchoring of a protein to a membrane concerns its sensitivity to bacterial PI-PLC^33^. Since exosomal membrane has the same topology as plasma membrane, MTf would be expected to be surface-accessible, as previously shown for other GPI-proteins (CD55, CD58 and CD59) in reticulocyte exosomes^14^. The rationale was thus to treat exosomes with PI-PLC to convert the membrane form to a soluble MTf released from the vesicle surface by cleavage of the diacylglycerol moiety from the GPI anchor (Figure 4A). Exosomes purified from SK-MEL-28 culture supernatant were thus incubated with PI-PLC from *Bacillus cereus* before re-pelleting the exosomes. In these conditions, MTf was no longer detected in the pelleted vesicles after PI-PLC treatment, but instead was retrieved in the centrifugation supernatant (Figure 4B). Note that a small amount of MTf was also found in the supernatant without PI-PLC treatment of exosomes. In contrast, neither the tetraspanin CD9 nor the cytosolic chaperone hsc70 were released from the treated vesicles, therefore confirming integrity of the vesicles.

**Figure 4:**
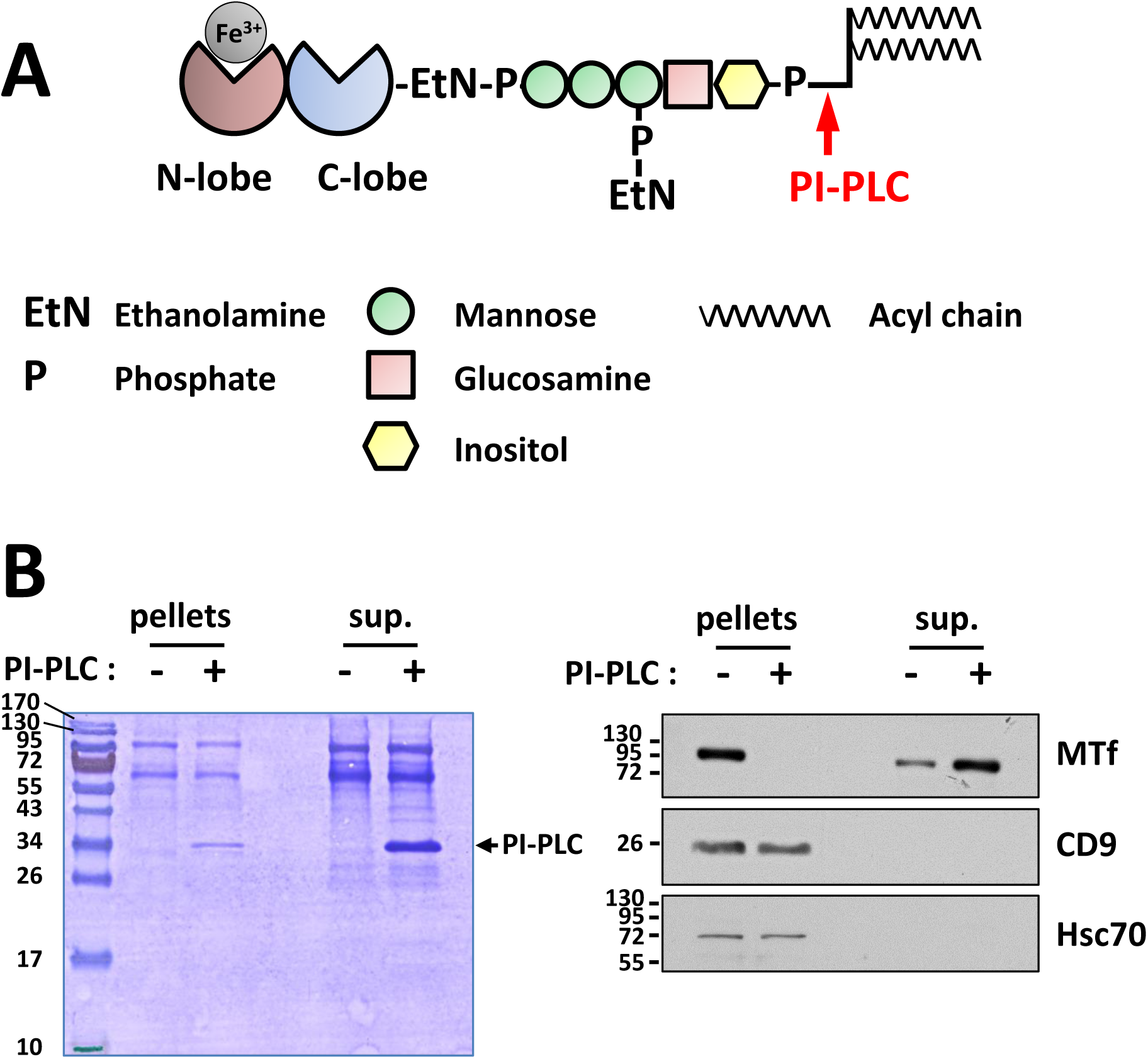
PI-PLC sensitivity of melanotransferrin. **A)** Melanotransferrin structure. MTf has 39% homology with serum transferrin. In contrast to serum transferrin, only the N-lobe of MTf is a Fe-binding site. MTf is bound to the cell membrane by a glycosylphosphatidylinositol (GPI) anchor. The specific cleavage by PI-PLC is indicated in red. **B)** Exosomes (5 µg) from SK-MEL-28 were incubated for 30 min at RT with or without PI-PLC from *Bacillus cereus* (1 u) before pelleting. Pellets and supernatants (sup.) were then loaded on SDS-PAGE and analyzed by Western blot for MTf and the indicated proteins.

Overall, these data demonstrated that melanotransferrin was efficiently sorted in exosomes – potentially through its glycolipid anchoring – and was surface-accessible.

### Melanotransferrin is specifically found in EVs from melanoma cells

Melanotransferrin was first described as a marker antigen for human melanoma cells^15^. MTf has actually also been detected in other neoplastic cells and fetal tissues, but only in small quantities in normal tissues^30,34^. We thus assessed the presence of melanotransferrin in vesicles secreted by other human cancer cell lines. For that, we used crude vesicle pellets – i.e. without sucrose gradient purification of the cell secreted vesicles – for a broad assessment of the MTf status. As shown in Figure 5A, melanoma cell lines released vesicles containing high amounts of MTf since, as already demonstrated, MTf was enriched in vesicle pellet compared to the same protein amount of corresponding cell lysates loaded on SDS-PAGE. Note also that, in these cell lines, conventional exosomal markers – including syntenin, Tsg101, CD63, Alix – were similarly enriched in the EV fractions, in contrast to other proteins (e.g. LDH-A, GAPDH) that were more abundant in cell lysate compared to the vesicular fraction. Breast adenocarcinoma MCF7 cells showed similar enrichment of conventional exosomal markers in EVs, but did not express MTf and consequently did not release vesicle-associated MTf. Colon cancer Caco2, chronic myelogenous leukemia K562 and cervix adenocarcinoma HeLa cells were tested and did not express/secrete vesicle-associated MTf. Note however that in these three cell lines, conventional exosomal marker enrichment was less marked in the vesicle pellets compared to cell lysates. For example, syntenin was found in slightly higher amounts in vesicles compared to cell lysates, while Tsg101 presented no difference between the two fractions. Similarly, CD63 tetraspanin was very faintly detected in these cell lines in our conditions, whereas the status of other proteins (TfR, flotillin) in EVs was highly dependent on the cell type. This could possibly be explained by the expression of these proteins by the cell lines, and/or more likely, by the relative number of endosomal-derived vesicles present in the crude 100 000 x g pellets. Figure 5B recaps, for the main proteins tested, the enrichment index – i.e. EV/cell lysate ratio – calculated for the different cell lines from Figure 5A. The enrichment index of classic exosomal markers – CD63, Tsg101, syntenin – was variable, but clearly noteworthy for melanoma cell lines and MCF7 cells (from 2.9 to 58.3). Similarly, MTf enrichment in EVs of melanoma cells was noteworthy (around two-fold) while, in contrast, the LDHA enrichment index was less than 1 in all of the tested cell lines.

**Figure 5:**
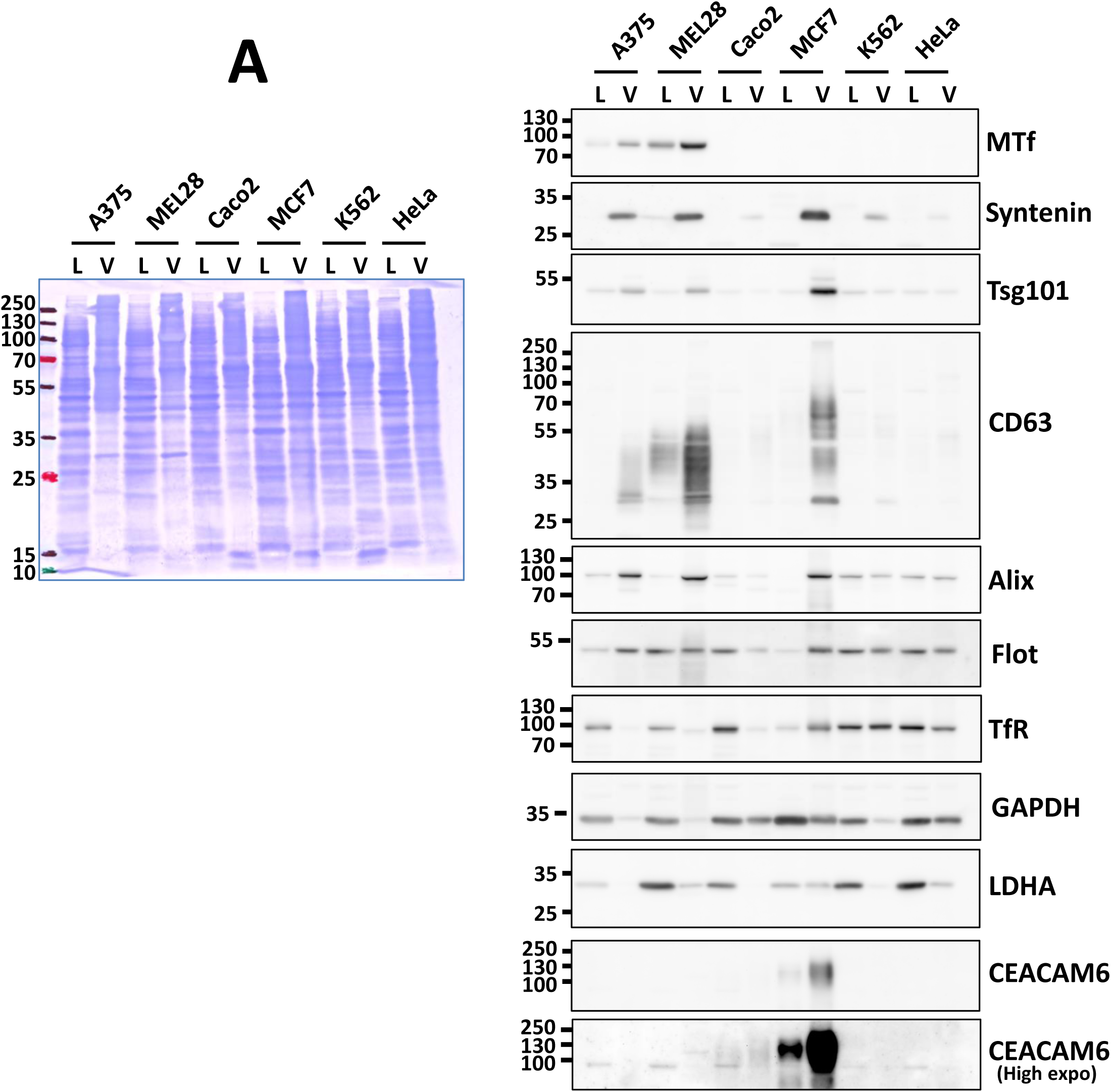
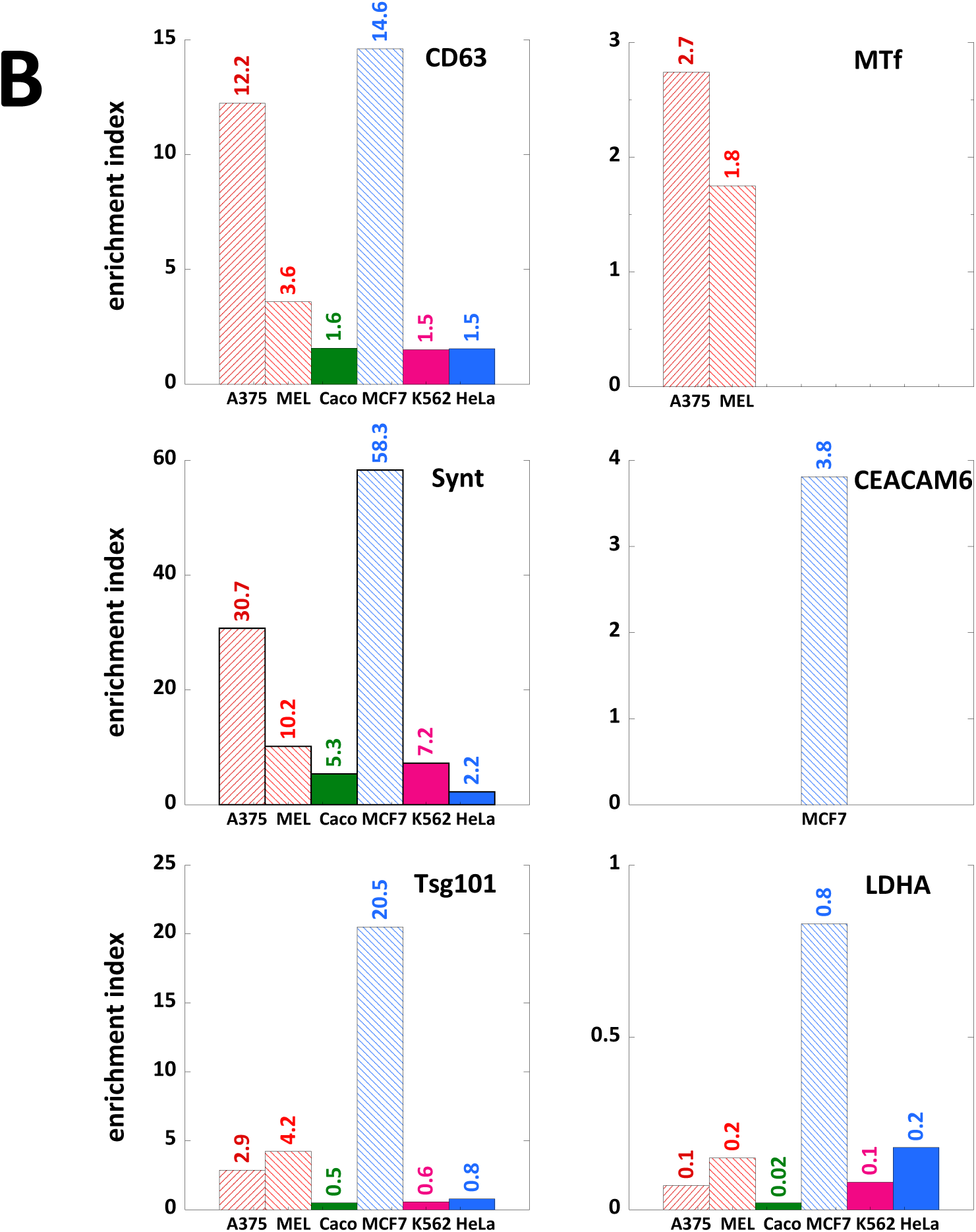
Melanotransferrin is specifically found on melanoma EVs. **A)** 5 µg protein of crude pellets of secreted EVs (V) and corresponding cell lysates (L) of A-375, SK-MEL-28, MCF7, Caco2, K562 and HeLa cells were loaded on 12% SDS-PAGE and analyzed for MTf and indicated proteins by Western blot. Coomassie staining of the PVDF membrane is shown (left panel). **B)** Enrichment index (EV/cell lysate ratio) of some proteins of interest – i.e. conventional exosomal markers, GPI-APs and a cytosolic protein – calculated for the different cell lines from Figure 5A.

To substantiate our assumption of lipid domain involvement in protein sorting, we assessed the presence of another GPI-anchored protein (GPI-AP) in these crude pellet fractions. CEACAM6 is a member of the carcinoembryonic antigen-related adhesion molecule (CEACAM) family, which is linked to the membrane through GPI linkage. CEACAM6 is associated with osteosarcoma metastasis^35^ and overexpressed in several cancers^36-38^. Very interestingly, CEACAM6 was substantially expressed in breast adenocarcinoma MCF7 cells, as reported in a previous study^39^, and strongly enriched in the EV fraction (3.8-fold) – compared to cell lysate (Figure 5B) – thus supporting our assumption of a lipid raft-related process. Note that with high exposure, CEACAM6 could also be detected in Caco2 cells with slight enrichment in EVs compared to cell lysate.

These data clearly showed that specific GPI-APs could be highly expressed in various cancer cell lines, and that they tended to accumulate in the membrane of secreted vesicles.

### MTf is associated with CD63-positive exosomes

To gain further insight into the association of MTf with the vesicles secreted by melanoma cells, we immunoisolated purified A-375 exosomes using magnetic beads coated with anti-CD63 antibody. As shown in Figure 6, exosomes were pelleted with beads only when coupled to CD63 antibodies, as indicated by the co-immunoisolation of exosomal markers such as Tsg101, Alix and hsc70. Note that these markers are cytosolic proteins proving the integrity of the immunoisolated vesicles. MTf was detected in the CD63-immunoisolated exosomes to a similar extent compared to conventional exosomal markers. Even though we did not isolate all the CD63-positive exosomes in our conditions, similar co-immunoisolation yields of exosomal markers – as well as our protein of interest – suggested that MTf was homogenously associated with CD63-positive vesicles. This prompted us to set up an assay to quantify exosomal MTf based on the presence of the two proteins – CD63 and MTf – on the surface of vesicles.

**Figure 6:**
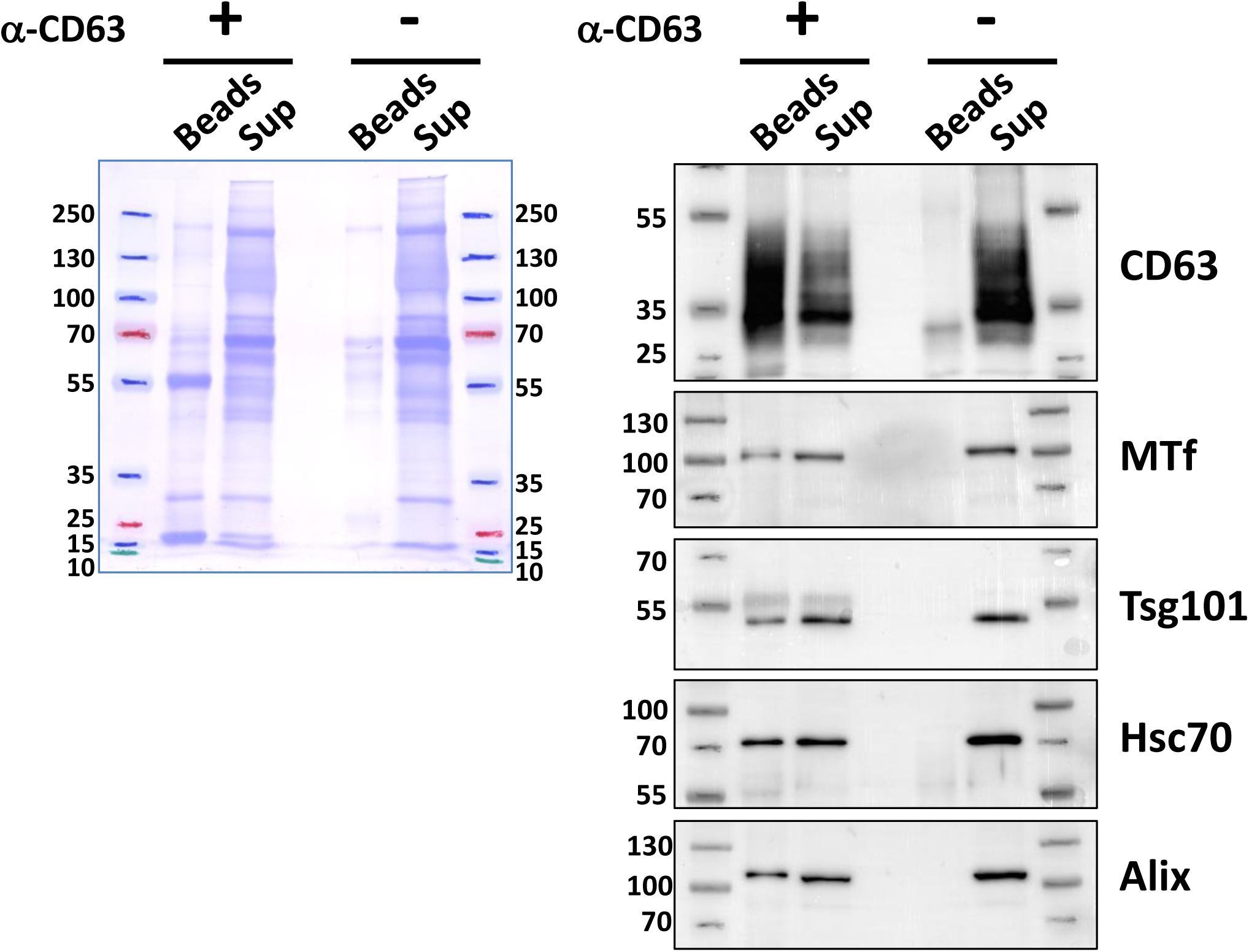
Melanotransferrin and CD63 are colocalized on melanoma exosomes. Immunoisolation of A-375 exosomes (25 µg) was carried out using Dynabeads™ protein G with or without anti-CD63 as described in the Materials and Methods. Washed beads (Beads) and half of the supernatants (Sup) were loaded on 12% SDS-PAGE and analyzed for the indicated proteins by Western blot (right panel). Coomassie staining of the PVDF membrane is shown (left panel).

### Exosomal MTf quantification by ELISA

An immunocapture-based ELISA was set up using rabbit anti-CD63 antibody to capture exosomes on the well surface and mouse anti-MTf antibody to detect exosomal melanotransferrin (Sup. Figure 1A). A first experiment (Figure 7A) using purified exosomes from the SK-MEL-28 cell line showed that the assay was functional, although the amounts of exosomes used were obviously oversized. However, note especially that for the highest exosome amount (approx. 8.3 µg protein), the absorbance measured at 450 nm was almost negligible when detergent was added (red dot) or when omitting the capture anti-CD63 antibody (blue dot), thus highlighting the relevance of our ELISA for measuring exosomal MTf through CD63 immobilization of the vesicles. We thus lowered the amount of exosomes assayed using a 1 ng to 1 µg protein range. Besides, we treated the highest amount of exosomes with PI-PLC as another way to determine the specificity of the assay. As shown in Figure 7B, 12 ng of protein was the minimum amount of purified exosomes for which exosomal MTf could be detected. Importantly, note that the corresponding absorbance threshold was very similar to that obtained when exosomes were treated by PI-PLC (red dot), to remove the exosomal-anchored MTf. As previously shown (Figures 1A, 1C, 2C and 5A), MTf was more expressed in SK-MEL-28 compared to A-375 cells and consequently in the respective secreted exosomes. For the sake of precision, we assessed ELISA quantitation of exosomal MTf in the two kinds of exosomes using the same number of vesicles – 10^9^ vesicles as determined by NTA – instead of the amount of protein. As shown in Figure 7C, exosomal expression of MTf was higher on vesicles from SK-MEL-28 cells compared to those released by A-375 cells in agreement with the cellular expression. We obtained similar results using another antibody (Abcam, ab201922) for MTf quantification (not shown).

**Figure 7:**
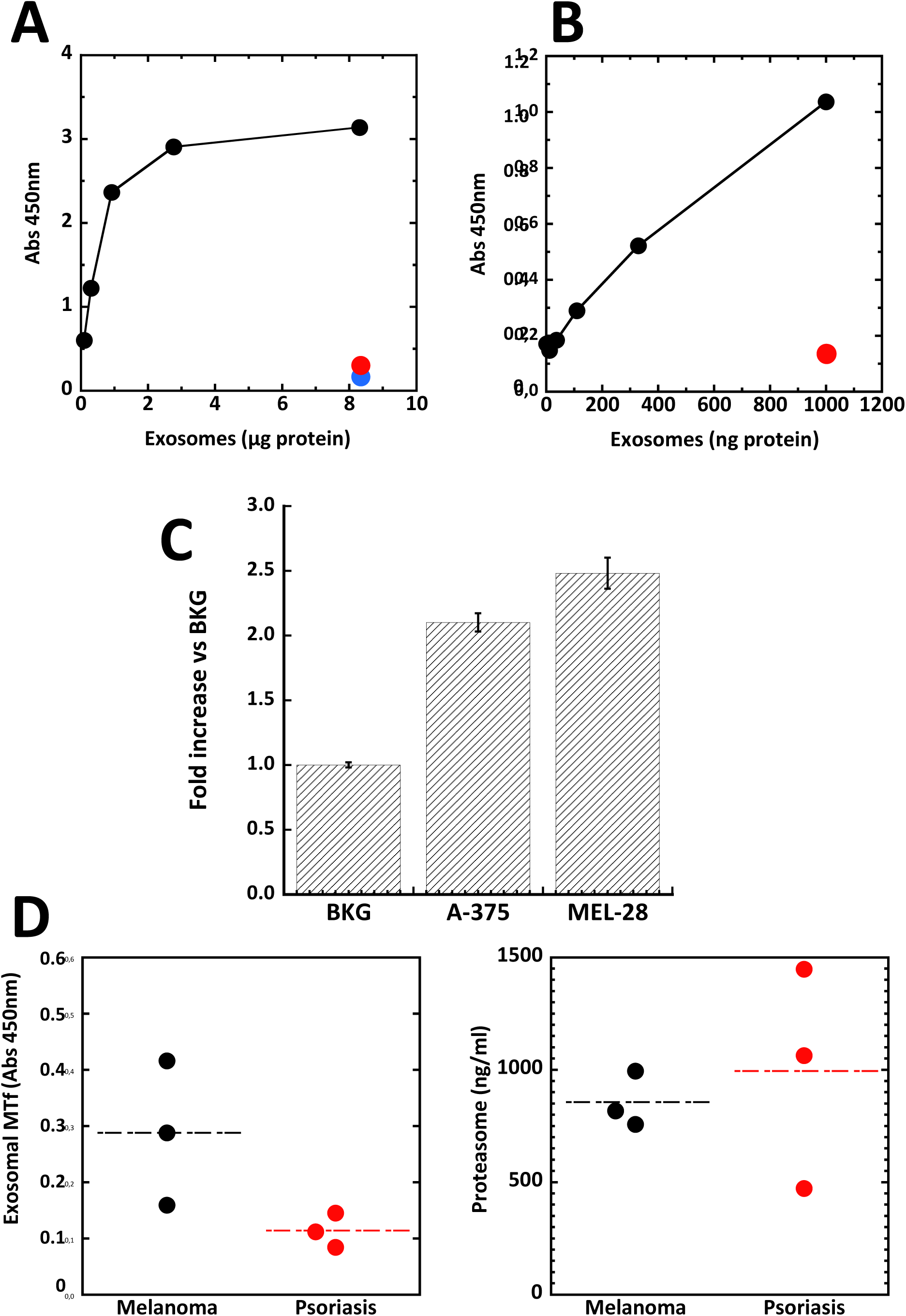
ELISA detection of exosomal melanotransferrin. **A)** Increasing amounts of purified SK-MEL-28 exosomes were adsorbed on wells coated with anti-CD63 antibodies (100 ng). MTf was revealed as described in the Materials and Methods using antibody (Ab201922) from Abcam. As controls, at the highest amount of exosomes (8.3 µg), vesicles were pre-treated with 1% Triton X-100 (red dot) or anti-CD63 antibody was omitted (blue dot). **B)** The lower range of purified exosomes was used to detect MTf as in (A). As a control, the highest amount of exosomes (1 µg) was pre-treated with 0.5 u PI-PLC (red dot) before ELISA readout. **C)** An equal number of exosomes (10^9^ vesicles) from A-375 and SK-MEL-28 cells were assayed by ELISA for MTf detection using antibody (MAB81751) from R&D. A typical experiment is presented (mean ± SD, n=3). Background (full assay in the absence of exosomes) was set to 1. **D)** Plasma of patients (three melanoma and three psoriatic) were used for exosomal MTf ELISA (left panel) and compared to p-proteasome ELISA (right panel). MTf was detected using Ab201922 (as in A and B). Mean plasmatic levels and optimal p-proteasome cutoff to diagnose metastatic melanoma was set at 387 ng/ml^5^.

We compared plasma from melanoma vs psoriatic patients to test our concept of exosomal MTf as melanoma biomarker. As already mentioned, plasma proteasome (p-proteasome) levels could be used to discriminate metastatic melanoma patients^5^. However, other diseases – even non-cancerous – can induce high p-proteasome levels, as especially shown for psoriasis patients with high p-proteasome levels^40^. Three plasma samples from non-cancerous psoriatic patients and three from melanoma patients (grade III-IV) having high plasma proteasome levels were tested for exosomal MTf using our ELISA. As shown in Figure 7D, the p-proteasome levels measured by quantitative ELISA were in a similar range for the two diseases (right panel). However, the two diseases were clearly distinguished through exosomal MTf quantitation (left panel), thus highlighting its relevance as a melanoma biomarker.

This study provided evidence in favour of the potential use of exosomal melanotransferrin as a melanoma biomarker. Yet we are aware that this finding would have to be confirmed and improved by assessing larger patient cohorts, determining its efficiency in detecting the various stages of metastatic melanoma progression and evaluating its capacity to assess therapeutic treatment, possibly using more sensitive detection approaches ^41,42^.

## Discussion

Exosomes are mainly studied for their capacity to transport membrane and cytosolic molecules from producer to recipient cells. As such, exosomes have been shown to change the behavior of recipient cells after uptake^43^. We are more interested in the biogenesis process of these vesicles^28^. Since the exosome composition is related to the producing cells, they could also be potential biomarkers during the course of diseases. The ideal situation would be to specifically detect exosomes released by diseased cells. This raises several problems related to the amount of analyzed exosomes, the presence of other kinds of vesicles/particles in biological fluids used for detection and the specificity of the exosomes considered amongst those secreted by cells unrelated to the disease. We were recently interested in exosomes and proteasomes released in the extracellular space by melanoma cell lines^18^. While levels of plasmatic proteasome (p-proteasome) were demonstrated to be correlated with metastatic melanoma progression^5^, there is no way to clearly identify the cells that release proteasome 20S particles in the plasma of patients. Moreover, increased levels of p-proteasome were detected with the progression of other types of cancer^44-46^ and inflammatory disease, such as psoriasis^40^. On the other hand, studies to find a signature for melanoma exosomes have been carried out using proteomic^22^ and miRNA^47^ analyses. These studies did not generate clear-cut information, and the various ‘specific’ molecules were mostly present in the cytosolic content of exosomes, thus rendering them less easy to detect.

We had previously shown that GPI-anchored proteins – i.e. CD55, CD58, CD59 – were secreted with reticulocyte exosomes via their association with raft domains^14,27^. Actually, other proteins – e.g. tetraspanins, flotillin – having affinity for lipid domains are regularly found in exosomes^8^. We thus hypothesized that MTf specifically overexpressed in melanoma cells could be sorted in exosomes through the same lipid raft-related process. As shown in Figures 2-4, MTf was indeed very efficiently secreted with exosomes from melanoma cell lines and was associated with the exosomal membrane through a PI-PLC-sensitive GPI-anchor. GPI-anchoring rules out any direct interaction of MTf with cytosolic proteins such as ESCRT proteins or syntenin. Instead, we found the presence of other lipid raft associated proteins (Lyn, flotillin), in agreement with the potential sorting of MTf in association with this lipid domain. Some exosomal proteins were recently demonstrated to have an inside-out topology^48^. This first set of experiments also revealed that the topology of MTf in the exosomal membrane was as expected, i.e. exposed on the vesicle surface and thus accessible for antibody recognition. We however noted (Figure 4B) that a small amount of MTf was recovered in the supernatant of exosomes untreated by PI-PLC. It is unclear if this corresponded to a described secreted soluble form of MTf^31^ or whether, more likely, it was released as a result of some experimental factor. Note also the high efficiency of GPI cleavage that could have been due to the high curvature of the exosomal membrane favoring accessibility/insertion of PI-PLC in the membrane^49^. Melanotransferrin was anchored in CD63 positive exosomes, as demonstrated by co-immunoisolation with CD63 (Figure 6).

We thus assessed the quantification of exosomal MTf by ELISA (Sup Figure 1A) and found that this assay was relevant (Figure 7A,B) and allowed detection of MTf using a scant amount of purified exosomes (Figure 7B). Moreover, the higher expression of MTf in SK-MEL-28 cells compared to A-375 cells – and consequently in the respective secreted exosomes (Figure 5) – coincided with the ELISA quantitation findings, as shown by analyzing an equivalent number of exosomes secreted by the two cell lines (Figure 7C). As final evidence of potential interest, measuring exosomal melanotransferrin in plasma from patients allowed us to discriminate between two diseases – melanoma and psoriasis – that otherwise have similar levels of p-proteasome, another potential biomarker for metastatic melanoma (Figure 7D).

Exosomal melanotransferrin might be a better prognostic biomarker of metastatic melanoma compared to LDH levels in blood since it is directly and specifically related to melanoma cells. Clinical use of this biomarker could especially be considered for personal monitoring of new melanoma treatments involving, for instance, small molecule inhibitors of BRAF and MEK, or immune checkpoint inhibitors. The use of lymphatic exudates, which were shown to have a higher exosome concentration compared to plasma of melanoma patients^50,51^, could also be considered. Moreover, MTf is present on the surface of exosomes, thus allowing easy detection using antibodies. Binding experiments with MTf antibodies on the surface of SK-MEL-28 cells estimated the presence of up to 380,000 sites/cell ^15,30^ even though MTf is not expressed at such high levels in all melanoma cell lines. Yet, as shown (Figures 2, 5), we noted clear sorting/enrichment of MTf in exosomes compared to cell lysates whatever the cellular expression levels, which might offset this expression peculiarity, especially when used for patient-specific follow-up. It might also be of interest to increase the sensitivity of exosomal melanotransferrin ELISA using new detection technologies^41,42^, or combining detection with another likely biomarker molecule on melanoma exosomes, such as GD3 ganglioside^52,53^. Conversely, other exosomal GPI-APs might be used for prognostic assessment of various cancers. As shown here, CEACAM6 was highly enriched in MCF7-secreted vesicles, while faintly present in Caco2-secreted vesicles and being virtually absent from the other tested vesicles. However, our data also revealed that such ELISA detection methods must also include the use of an accurate prey – presently CD63 – for exosome capture, which could depend on the vesicle-producing cells. At this point, note that BRAF mutation (V600E) – a critical mutation in melanoma – has been shown to correlate with overexpression of CD63^54^.

Finally, MTf association with melanoma exosomes could also be used as a research tool to specifically purify these vesicles and to look for other kinds of specific exosomal markers (e.g. miRNAs).

The cellular function of melanotransferrin is still not very clear. Because of its homology with transferrin, a role in iron metabolism was first suggested^55^ before it was demonstrated that MTf did not provide Fe to melanoma cells^56^. More recently, various roles of MTf have been reported in relation with cancer progression^24^, such as cell invasion^57^, melanoma tumorigenesis^58^ and angiogenesis^59^. The presence of MTf on the surface of melanoma exosomes might shed light on these processes. For example, it has been shown that melanoma exosomes contribute to pre-metastatic niche settlement^60^. Remodeling the extracellular matrix is one of the events involved in this process. In agreement, by zymography^61^ we found that exosomes secreted by both A-375 and SK-MEL-28 cell lines hold gelatinase activities (Sup. Figure 2A), which might correspond to matrix metalloproteinases MMP-2 and MMP-9^62^. Note that these gelatinase activities were not detected using equivalent protein amounts of cell lysates. Moreover, the urokinase-type plasminogen activator receptor (uPAR) is a GPI-anchored protein that has been demonstrated to promote melanoma cell migration and invasion^63^. One might speculate that uPAR – secreted with melanoma exosomes^22^ – is also associated with lipid rafts on the membrane of melanoma exosomes which could thus behave as a pro-degradation platform. Actually, GPI-anchored proteins MTf and uPAR respectively bind to plasminogen and urokinase^64,65^, and could locally favor plasmin production. This plasmin production might in turn participate in matrix metalloproteinase activation (Sup. Figure 2B). In this case, exosomes could merely bind to the selected microenvironment without the need of fusing with any cell membrane. Moreover, molecular transfer of GPI-proteins from one membrane to another has been reported both *in vitro*^66^ and *in vivo*^67,68^ and, if this occurs with MTf and uPAR, could sustainably be involved in microenvironment remodeling.

Finally, it is also interesting to note that MTf has been used as carrier to target various compounds across the blood brain barrier (BBB)^69,70^, and that melanoma brain metastasis is inhibited by targeting melanotransferrin at the cell surface^71^. Moreover, melanoma exosomes have been shown to cross the BBB and possibly to participate to BBB defects^72^. Yet it remains to be determined whether melanoma exosomes can cross/damage the blood brain barrier using MTf.

## Acknowledgments

This work was supported by CNRS, the University of Montpellier and grants from the Cancéropôle *GSO* (MelanoProtEx) and the *Ligue Nationale Contre le Cancer - région Languedoc-Roussillon*. We warmly thank Dr. Laure Yatime (LPHI, Univ. Montpellier) and Dr. Stéphane Bodin (CRBM, Univ. Montpellier) for sharing reagents. We acknowledge the imaging facility MRI, member of the national infrastructure France-BioImaging infrastructure supported by the French National Research Agency (ANR-10-INBS-04, «Investments for the future»).

## Supporting Information

**Supplementary Figure 1: Immunocapture-based ELISA diagram of exosomal MTf quantification on CD63-positive exosomes**.

**Supplementary Figure 2: Potential involvement of melanotransferrin in MMP activation. A)** Exosomes and cell lysates (10 µg) from A-375 and SK-MEL-28 cell lines were assayed for gelatin zymography as described in the Materials and Methods. White bands correspond to degradation of gelatin impregnated in the gel. MMP-2 and MMP-9 are two matrix metalloproteinases with gelatinase activity. **B)** Scheme representing the possible contribution of exosomal MTf and uPAR on MMP activation.

