## Supplementary figures and images for "Exosomal melanotransferrin as a potential biomarker for metastatic melanoma"

### supplemental figure 1

#### Slide 1
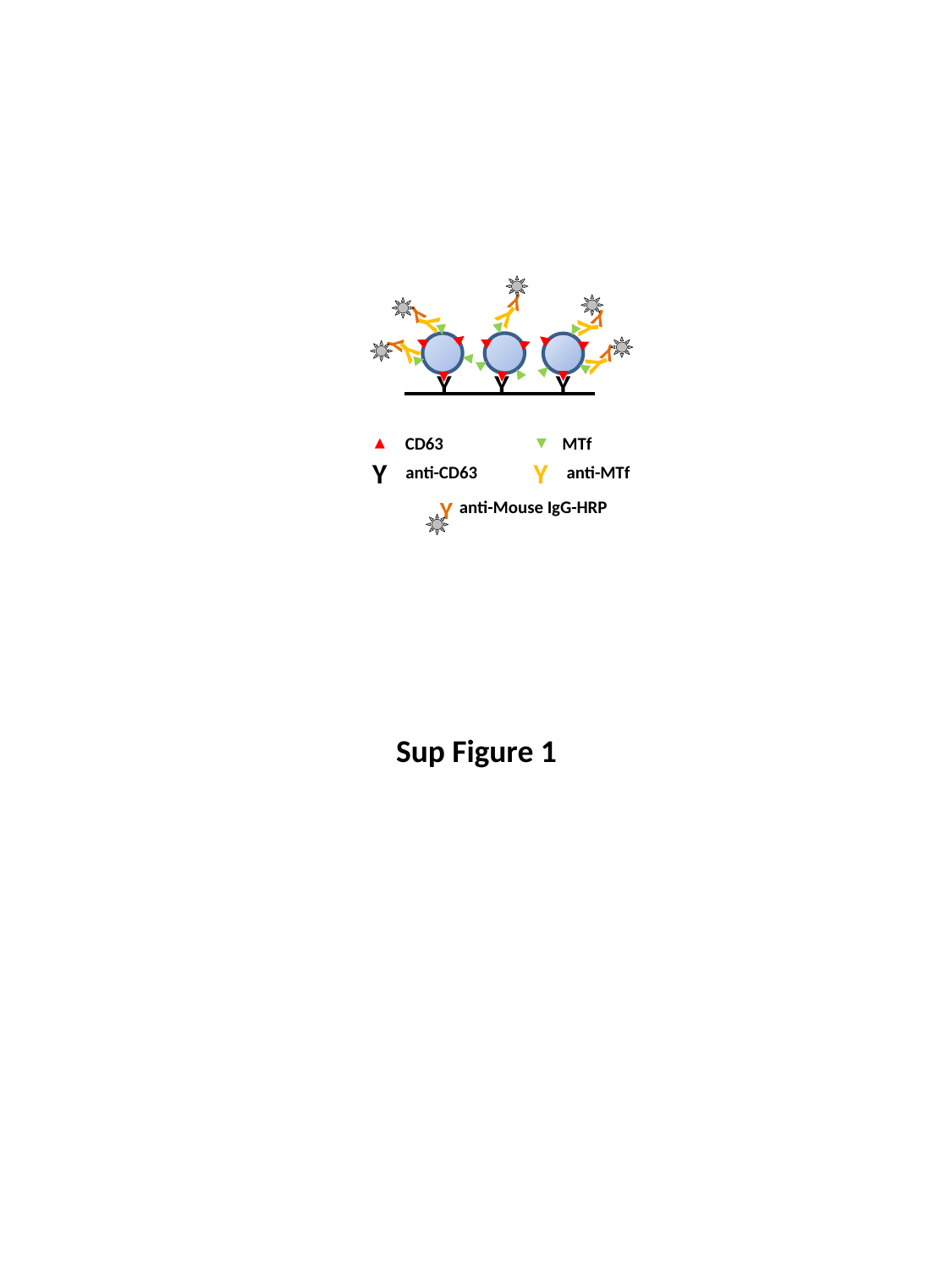

Y
Y
Y
Y
Y
Y
Y
Y
Y
Y
Y
Y
Y
CD63
MTf
Y
Y
anti-CD63
anti-MTf
Y
anti-Mouse IgG-HRP
Sup Figure 1
