## supplemental figure 2 for "Exosomal melanotransferrin as a potential biomarker for metastatic melanoma"

### Slide 1
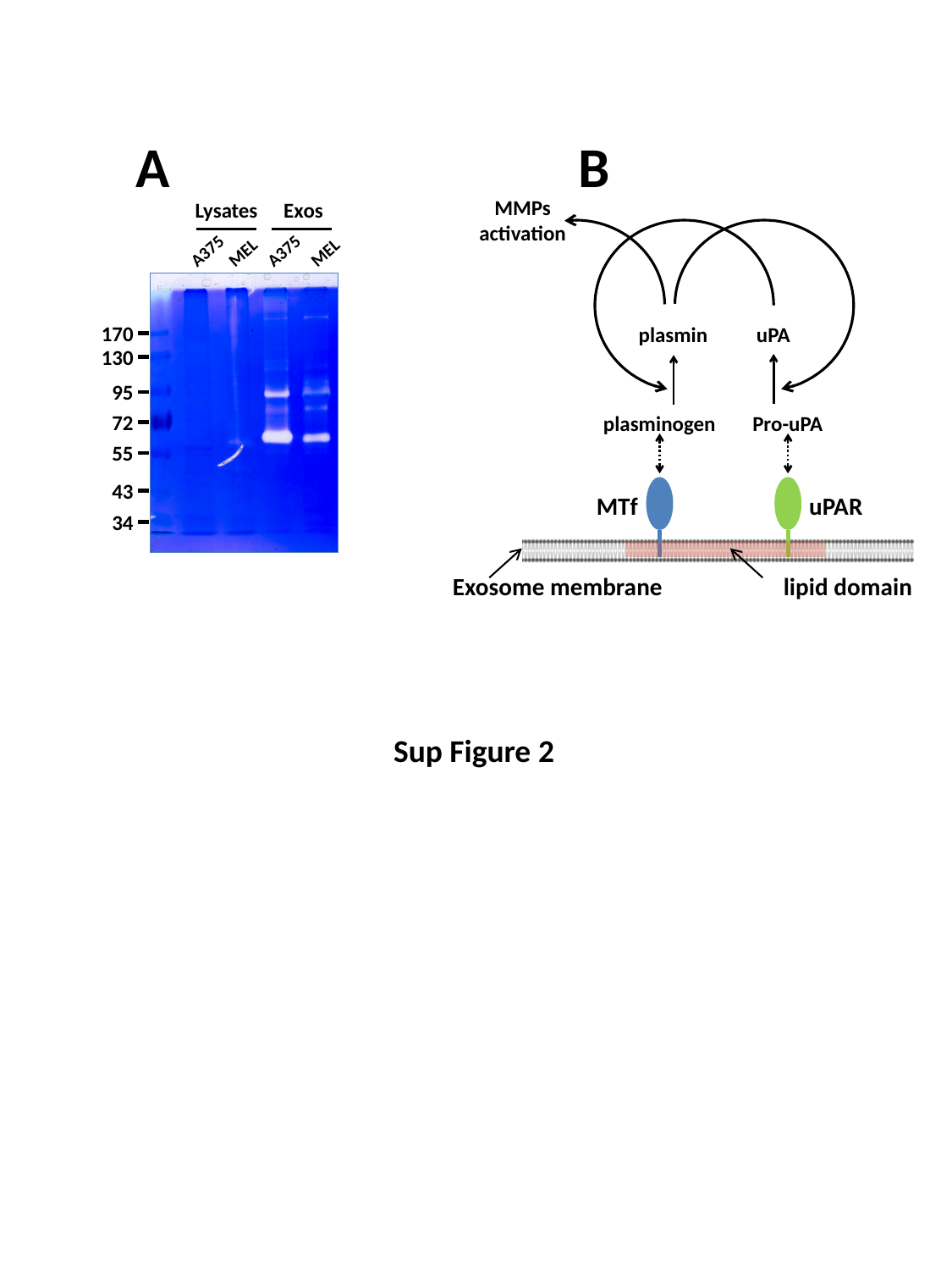

A
B
MMPs
activation
plasmin
uPA
plasminogen
Pro-uPA
MTf
uPAR
Exosome membrane
lipid domain
Lysates
Exos
A375
A375
MEL
MEL
170
130
95
72
55
43
34
Sup Figure 2
